# Additive Effects of Sleep Loss, Psychological Distress and Physical Inactivity on Cognitive Failures in Young Adults

**DOI:** 10.64898/2026.06.11.731711

**Authors:** Anushka Sarkar

**Affiliations:** Neuroscience Unit, Jawaharlal Nehru Centre for Advanced Scientific Research, Bangalore 560064, India

**Author notes:** Donders Institute for Brain, Cognition and Behaviour, Radboud University, Nijmegen, Netherlands.

**Keywords:** cognitive burden, sleep restriction, physical activity, psychological distress, young adults

## Abstract

Young adults frequently report cognitive complaints often attributed to sleep loss alone. However, subjective cognitive functioning is shaped by broader lifestyle and affective factors. Cross-sectional data were analyzed from 530 young adults (mean age 22.1 *±* 2.3 years) to examine the independent, interactive, and cumulative associations of short sleep duration, low physical activity, and psychological distress with everyday cognitive failures. Cognitive failures were strongly associated with sleep duration, physical activity, sleep quality, and distress in univariate analyses. However, hierarchical regression revealed that psychological distress, poor sleep quality, and short sleep duration were the dominant independent correlates of cognitive failures, collectively explaining a substantial proportion of variance in Cognitive Failures Questionnaire scores (*R*^2^ = 0.585, *p <* 0.001). In contrast, the apparent protective effect of physical activity was not observed after adjustment for sleep and distress (*p* = 0.976), and no significant sleep-by-physical activity interaction was observed. Further, cumulative risk modeling demonstrated a robust dose-dependent relationship, with cognitive failures increasing progressively as behavioral and psychological risk factors accumulated (*p*_trend_ *<* 0.001). Individuals exposed to all three risk factors exhibited more than double the cognitive failure burden observed in individuals with no risk factors. These results indicate that the cognitive burden in young adults can best be described by a additive increase of behavioral and psychological risk factors as a function of the co-occurrence, rather than by the presence of compensatory effects of lifestyle risk factors. Interventions aimed at preserving cognitive function may therefore benefit from simultaneously targeting sleep health and psychological well-being rather than relying on physical activity alone to offset cognitive burden.

## 1 Introduction

Young adult populations, particularly college students [1], frequently experience subjective complaints of brain fog, memory lapses, and attentional deficits due to the combination of high academic pressure and irregular lifestyle patterns. These executive deficiencies are usually attributed by the classic biological paradigm to chronic sleep restriction, a hallmark of student life widely assumed to degrade prefrontal cortical function [3]. Yet, this linear, single-variable causal model is increasingly challenged by emerging epidemiological evidence.

Today, short sleep duration, high sedentary behavior, and elevated psychological distress frequently co-occur, posing a complex, multifaceted danger to everyday cognitive function. Controlling for lifestyle and affective factors in multivariable analyses often significantly weakens the independent relationship between sleep duration and cognitive decline. This suggests that emotional distress and physical inactivity may be misattributed to sleep-related cognitive failures [4, 5]. They are generally minor but frequent lapses in everyday memory, perception, and motor function, and serve as an early behavioral manifestation of disrupted cognitive maintenance. These subjective complaints are closely linked to emotional mood and perceived allostatic stress as they represent an individual’s lived experiences [6].

Physical activity (PA) has historically been theorized to serve as a neuroprotective moderator against these disruptions, with proposed mechanisms including the upregulation of Brain-Derived Neurotrophic Factor (BDNF) and enhanced systemic vascular health [7]. Some experimental evidence also suggest that exercise may preserve vigilance via affective and metabolic regulation, buffering the cognitive deficits induced by homeostatic sleep pressure [8]. On the other hand, sedentary behavior has been associated with elevated symptoms of depression and anxiety, independent of the mere absence of exercise [9]. Similarly, exercise-induced improvements in cognitive performance have been widely suggested across experimental paradigms [10]. But nevertheless, it remains fundamentally unclear whether physical activity serves as a true physiological buffer against the cognitive deficits induced by short sleep (a multiplicative effect), or whether sleep deprivation, inactivity, and distress merely accumulate to erode cognitive capacity (an additive effect).

This study aimed to formally quantify the independent, interactive, and cumulative role of sleep duration, physical activity, and psychological distress on daily cognitive failures. By employing rigorous hierarchical regression modeling, false discovery rate (FDR) corrections, and pre-specified sensitivity analyses, I hypothe-sized that rather than a simple multiplicative interaction, individuals presenting with an aggregated high-risk behavioral phenotype would exhibit significantly higher rates of cognitive failure in a strict, dose-dependent, additive manner.

## 2 Methods

### 2.1 Participants and Procedure

Participants were recruited through online dissemination of a survey link via university mailing lists, student networks, and social media platforms. A total of 530 young adults completed the survey (mean age 22.1 *±* 2.3 years; 55.3% female). The study primarily targeted college-going individuals to investigate lifestyle and psychological correlates of everyday cognitive functioning in a non-clinical population. Participation was voluntary and anonymous, and collected no personally identifiable information. Individuals reporting diagnosed neurological disorders or current use of sleep-modulating medications were excluded to minimize potential clinical confounds. All participants provided informed digital consent prior to participation, and study procedures were conducted in accordance with the Declaration of Helsinki.

### 2.2 Measures

#### 2.2.1 Cognitive Failures Questionnaire (CFQ)

Perceived Cognitive Difficulties (PCD) were assessed using a 9-item adapted version of the Cognitive Failures Questionnaire (CFQ) [11], specifically selected to capture attentional lapses and memory failures in daily functioning. Questions were drawn from the original 25-item CFQ and retained on the basis of factor loadings from prior psychometric work identifying a unidimensional attentional-failure factor [6, 11]. Internal reliability in the present sample was acceptable (Cronbach’s *α* = 0.78). Responses were summed to generate a continuous cognitive-failure score, with higher values indicating a greater frequency of subjective cognitive failures. Given the established conceptual overlap between self-reported cognitive failures and affective state [6], CFQ scores were interpreted as an index of subjective cognitive concern and everyday functional lapses rather than objective cognitive impairment.

#### 2.2.2 Sleep Metrics

Self-reported habitual sleep duration was evaluated both as a continuous variable (hours) and dichotomized into short sleep (*<* 7 h) versus adequate sleep (*≥* 7 h), following consensus clinical guidelines [2]. Sleep quality was objectively scored utilizing the Pittsburgh Sleep Quality Index (PSQI), a validated self-report measure. Further, sensitivity analyses additionally modeled sleep duration as a three-level ordinal factor (*<* 6 h, 6–7 h, *≥* 7 h) to confirm the robustness of the dose-response trajectory.

#### 2.2.3 Physical Activity and Sedentary Behavior

Physical activity was quantified using metabolic equivalent (MET)-minutes per week via the short form of the International Physical Activity Questionnaire (IPAQ-SF) [12]. Participants were categorized into active (*≥* 600 MET-min/wk) and low PA/sedentary (*<* 600 MET-min/wk) groups based on standardized thresholds. Daily sedentary hours were additionally recorded as an independent continuous metric for lifestyle burden.

#### 2.2.4 Psychological Distress

Mental health status was evaluated using the 4-item Patient Health Questionnaire for Depression and Anxiety (PHQ-4) [13, 14]. A threshold score of *≥* 3 was used to categorize participants into a high-distress group for cumulative risk modeling, while the continuous score was used in regression frameworks to maximize statistical sensitivity to the affective signal.

### 2.3 Statistical Analysis

Analyses were conducted in Python 3.12.3 using the pandas, numpy, scipy, statsmodels, scikit-learn, matplotlib, and seaborn libraries. Prior to regression modeling, residuals from the full model were examined for normality using the Shapiro–Wilk test (W = 0.997, *p* = 0.610). A minor deviation in homoscedasticity was detected using the Breusch–Pagan test (LM = 16.00, *p* = 0.042), but this was deemed statistically acceptable given the sample size (*N* = 530). Multicollinearity was evaluated using variance inflation factors (VIF); all variables demonstrated acceptable independence (maximum VIF = 3.91 for the interaction term, with all main effects *<* 3.01). Bivariate correlations (Pearson’s *r*) were calculated to assess preliminary relationships. To control for type I error accumulation, all secondary predictor *p*-values were adjusted using the Benjamini–Hochberg false discovery rate (FDR) [16]. Groupwise mean differences in CFQ were evaluated using independent-samples *t*-tests and Cohen’s *d* effect sizes. To systematically isolate the predictors of cognitive failures, I constructed a four-block hierarchical ordinary least squares (OLS) regression model [15]. Variables were entered sequentially as follows: Block 1, demographics (age, gender); Block 2, lifestyle parameters (short sleep, active physical activity status, sedentary hours, PSQI); Block 3, psychological distress; and Block 4, the sleep *×* physical activity interaction. Since interaction testing carries an elevated risk of type I error, the term was subjected to a Bonferroni-corrected significance threshold (*α* = 0.0083). Finally, an additive cumulative risk analysis was conducted by summing the presence of the three primary binary risk factors (short sleep, low physical activity, and high distress) and fitting a linear regression trend.

## 3 Results

### 3.1 Prevalence and Bivariate Associations

The cohort exhibited a mean CFQ score of 7.83 *±* 3.04 (range 0.0–15.9). Behavioral risk factors were highly prevalent: 50.2% (*n* = 266) reported short sleep, 35.8% (*n* = 190) met criteria for low physical activity, and 34.2% (*n* = 181) exhibited high psychological distress. The mean total physical activity was 882 *±* 553 MET-min/wk, while the mean global PSQI score was 6.74 *±* 2.68, indicating widespread decline in sleep quality.

Bivariate correlation matrices revealed robust zero-order associations between the variables of interest and CFQ scores (Figure 1b). Higher cognitive failures were significantly correlated with increased psychological distress (*r* = 0.406, *p*_FDR_ = 3.21 *×* 10^*−*22^), poorer global sleep quality via PSQI (*r* = 0.567, *p*_FDR_ = 1.80 *×* 10^*−*45^), and greater daily sedentary hours (*r* = 0.122, *p*_FDR_ = 0.006). Conversely, CFQ scores were strongly inversely related to continuous sleep hours (*r* = *−*0.449, *p*_FDR_ = 2.94 *×* 10^*−*27^; Figure 1f) and total physical activity MET-minutes (*r* = *−*0.467, *p*_FDR_ = 1.69 *×* 10^*−*29^). Age (*p*_FDR_ = 0.297) and smartphone usage hours (*p*_FDR_ = 0.110) were not significantly correlated with cognitive failures.

**Figure 1:**
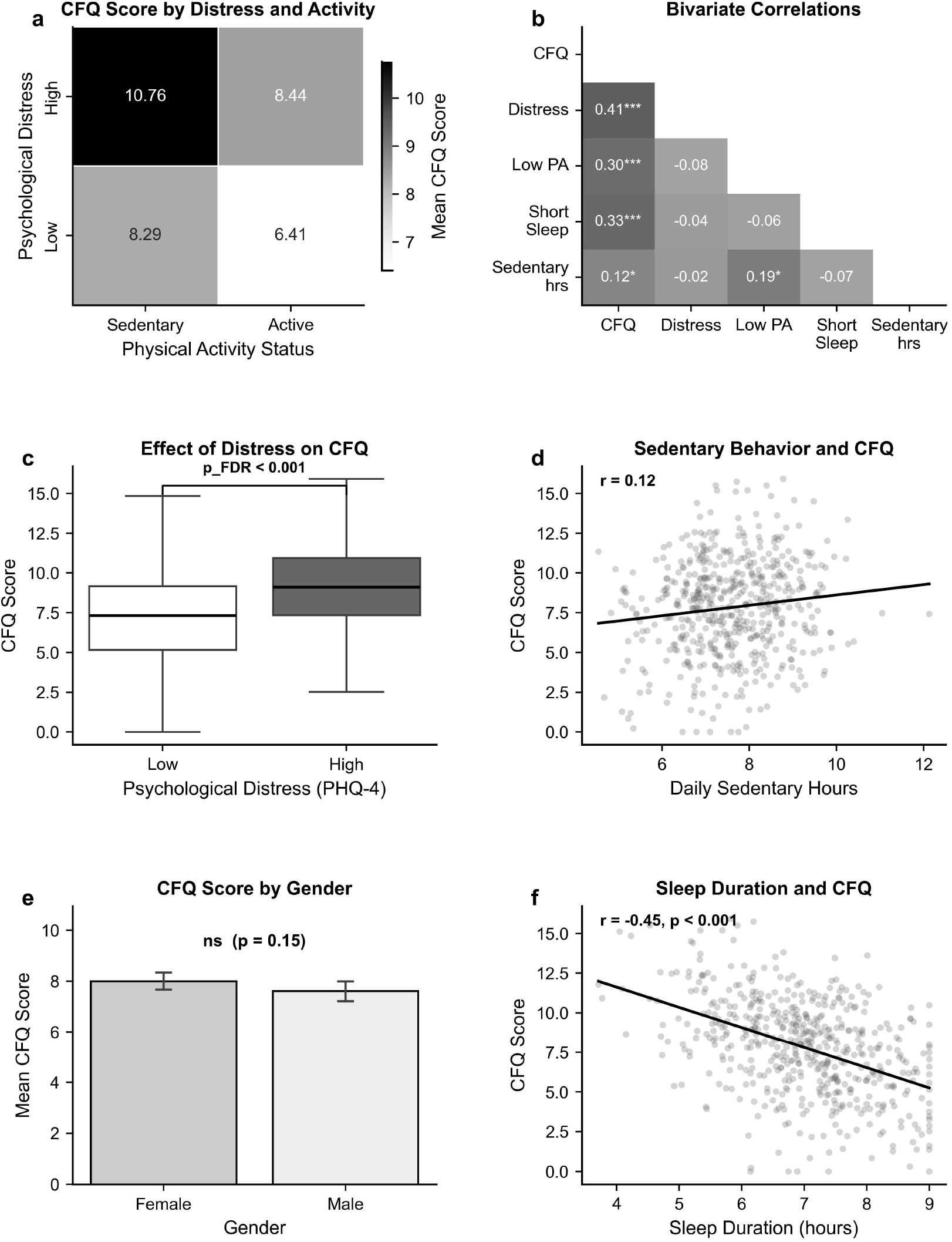
Epidemiological distribution and bivariate determinants of CFQ. (**a**) Joint density heatmap of CFQ stratified by physical activity status and distress level. (**b**) Bivariate correlation matrix of key risk factors demonstrating strong independent associations between CFQ, distress, sleep, and physical activity. (**c**) Psychological distress exerts a substantial influence on cognitive failures, with high-distress individuals reporting significantly elevated scores (*p <* 0.001). (**d**) Daily sedentary hours show a weak but significant positive correlation with CFQ. (**e**) No significant differences in CFQ scores were observed across genders (*p* = 0.153). (**f**) Sleep duration demonstrates a strong inverse relationship with cognitive failures (*r* = *−*0.45, *p <* 0.001).

### 3.2 Group Differences and Behavioral Phenotypes

Independent samples *t*-tests corroborated the bivariate findings across categorical groupings. Individuals exhibiting high psychological distress reported significant higher CFQ scores (9.18 *±* 0.20) compared to those with low distress (7.12 *±* 0.16; *t*(528) = 7.824, *p <* 0.001; Figure 1c). Similarly, the short sleep cohort reported significantly more cognitive failures than the adequate sleep cohort (8.84 *±* 0.18 vs. 6.81 *±* 0.17; *t*(528) = 8.150, *p <* 0.001, Cohen’s *d* = 0.668). A comparable effect magnitude was observed in the low PA group (9.05 *±* 0.20) relative to active individuals (7.14 *±* 0.16; *t*(528) = 7.237, *p <* 0.001, *d* = 0.626). Consistent with correlation findings, no significant gender differences were observed (Female: 8.00 *±* 0.18 vs. Male: 7.62 *±* 0.20; *t*(528) = 1.432, *p* = 0.153; Figure 1e).

When participants were stratified into integrated behavioral phenotypes (Figure 2b), individuals showing the “High Risk” profile (sedentary + short sleep; *n* = 88) reported the highest cognitive burden (10.33 *±* 0.28). This was significantly higher than the “Buffered” profile (active + short sleep; *n* = 178, mean = 8.10 *±* 0.21; *t* = 6.186, *p <* 0.001, Cohen’s *d* = 0.734).

**Figure 2:**
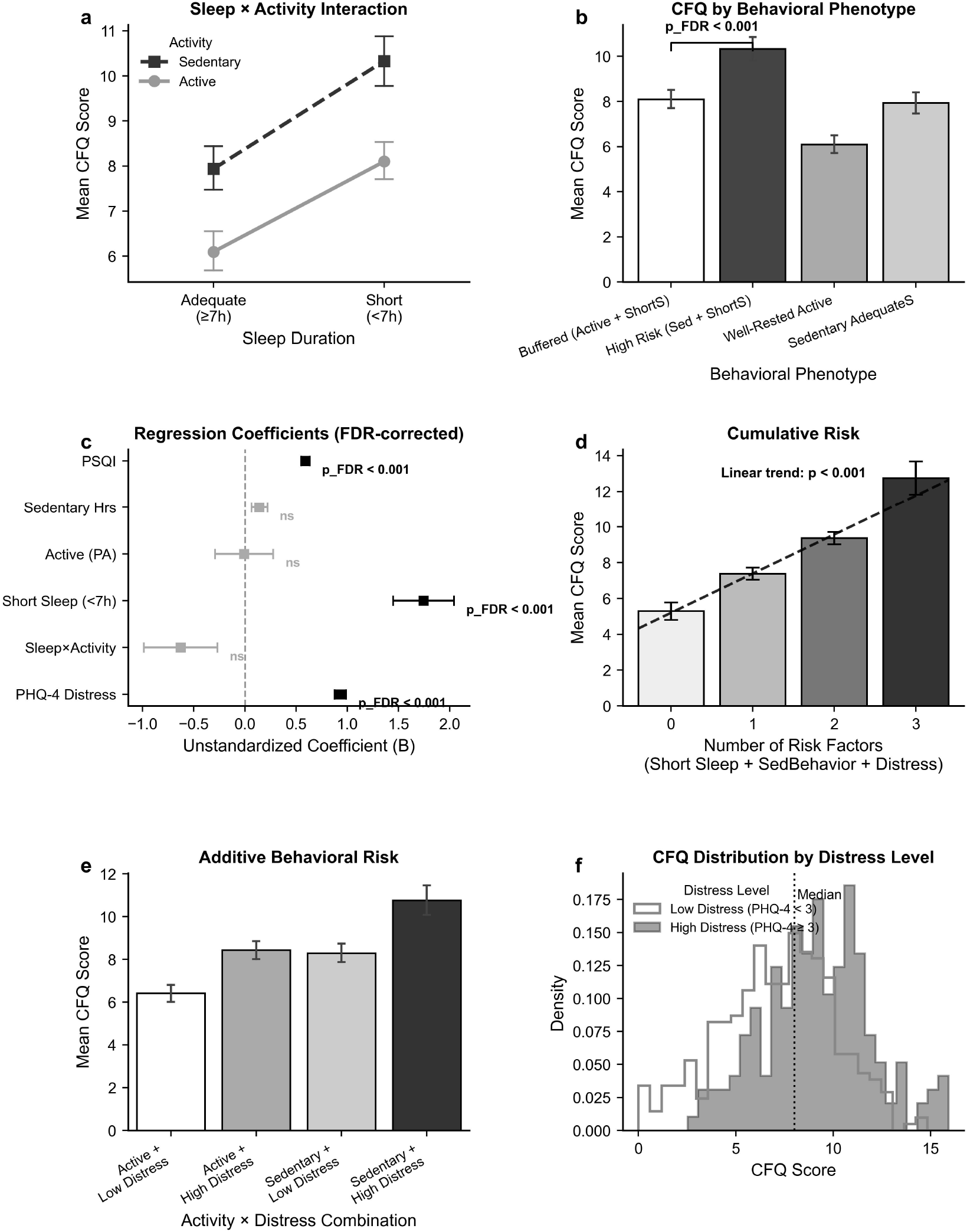
Statistical modeling of additive and interactive cognitive risk. (**a**) The Sleep × Physical Activity interaction shows a non-significant buffering trend (*p >* 0.05 after correction), indicating PA does not statistically eliminate the risk of short sleep. (**b**) Behavioral phenotypes demonstrate graded cognitive outcomes. (**c**) Forest plot of FDR-corrected regression coefficients. Psychological distress and PSQI dominate the model variance; the main effect of Physical Activity is completely nullified (*p* = 0.976). (**d**) Cumulative risk modeling demonstrates a strict linear dose-response escalation (*p*_trend_ *<* 0.001) in CFQ scores as the number of risk factors increases from 0 to 3. (**e**) Additive behavioral risk highlights that combining sedentary behavior with high distress yields the most severe cognitive impairments. (**f**) High psychological distress shifts the population CFQ distribution rightward, substantially increasing the density of severe cognitive complaints.

### 3.3 Hierarchical Regression Modeling

To disentangle these overlapping phenotypes and control for shared variance, a hierarchical regression was performed. The final multivariable model (Block 4) was highly robust, accounting for 58.5% of the total variance in CFQ scores (*R*^2^ = 0.585, *F* (8, 521) = 91.66, *p* = 2.45 *×* 10^*−*94^). Stepwise integration elucidated the underlying behavioral hierarchy. Demographic factors (Block 1) explained minimal variance (*R*^2^ = 0.005). The integration of lifestyle factors (Block 2) improved the model fit (Δ*R*^2^ = 0.360, *F*_change_ = 74.42, *p <* 0.001). Crucially, the addition of psychological distress (Block 3) captured a massive proportion of unique variance (Δ*R*^2^ = 0.216, *F*_change_ = 269.71, *p <* 0.001). The inclusion of the Sleep *×* PA interaction term (Block 4) provided only a marginal, non-significant improvement (Δ*R*^2^ = 0.002, *F*_change_ = 3.06, *p* = 0.080). In the final model (Figure 2c), short sleep (*B* = 1.744, *p*_FDR_ *<* 0.001, partial *η*^2^ = 0.062), global PSQI (*B* = 0.590, *p*_FDR_ *<* 0.001), and psychological distress (*B* = 0.931, *p*_FDR_ *<* 0.001) emerged as the dominant independent predictors. Distress solely accounted for over 34% of the variance in CFQ scores (partial *η*^2^ = 0.344), establishing it as the primary driver of perceived cognitive failures. Strikingly, the main effect of physical activity - which appeared robust in unadjusted bivariate testing - was entirely nullified upon controlling for sleep quality and distress (*B* = *−*0.008, *p* = 0.976). Furthermore, the interaction between short sleep and physical activity was not statistically significant following Bonferroni correction for multiple comparisons (*B* = *−*0.629, *p* = 0.081).

### 3.4 Sensitivity and Cumulative Risk Analyses

To validate the sleep architecture findings, sensitivity analyses were conducted using alternative variable constraints. Evaluating sleep as a continuous variable yielded consistent predictive results (*B* = *−*0.880, *p* = 2.10 *×* 10^*−*22^). Reparameterizing sleep into a 3-level ordinal scale confirmed a strict dose-response trajectory: compared to individuals sleeping *<* 6 hours, those achieving 6–7 hours (*B* = *−*1.181, *p* = 3.85 *×* 10^*−*6^) and *≥* 7 hours (*B* = *−*2.150, *p* = 1.51 *×* 10^*−*16^) exhibited significantly lower CFQ scores. Interestingly, applying a conservative reclassification to physical activity thresholds (+20%) yielded a nominally significant interaction (*B* = *−*0.846, *p* = 0.014), but it did not survive the pre-specified multiple-comparison threshold and should therefore be interpreted cautiously, and thus it remains secondary to the main additive effects.

Given the absence of a robust multiplicative interaction in the primary model, I evaluated the additive burden of the primary variables (Figure 2d). Cumulative risk modeling revealed a highly significant, perfectly monotonic linear trend (slope = 2.187, *r* = 0.578, *p* = 1.33 *×* 10^*−*48^). Participants with zero risk factors established the baseline cognitive failure threshold (5.30 *±* 0.25). The addition of each subsequent risk factor escalated the cognitive burden sequentially: one factor (7.39 *±* 0.17), two factors (9.37 *±* 0.17), culminating in severe cognitive lapses for individuals enduring all three risk parameters simultaneously (12.75 *±* 0.47).

## 4 Discussion

This work demonstrates that subjective cognitive failures in young adults are driven predominantly by an additive accumulation of lifestyle and psychological vulnerabilities, accounting for nearly 60% of the variance in CFQ scores. In a rigorous multivariable context, the cognitive impact of physical activity is largely subsumed by the pronounced consequences of disrupted sleep architecture and mental health status. Psychological distress alone uniquely explained over a third of the variance in cognitive failures (partial *η*^2^ = 0.344). While the CFQ’s nature as a self-report measure inherently overlaps with affective states [6], this covariance highlights how deeply shared the neural substrates of emotional regulation and cognitive tracking truly are. This psychological tax operates alongside the non-negotiable demands of sleep-dependent neural restoration, captured by the robust main effects of short sleep (*B* = 1.744) and poor sleep quality (*B* = 0.590). These findings align with systemic models of allostatic load, where chronic hyperarousal saturates executive capacity, leaving fewer neural resources for routine monitoring and memory retrieval [5, 17].Crucially, the data reveal a strict biological hierarchy rather than a multiplicative interaction: physical activity does not statistically buffer the deficits caused by short sleep once distress and sleep quality are controlled. While exercise promotes vascular and metabolic health [7], it cannot substitute for the precise synaptic downscaling and glymphatic clearance that occurs strictly during slow-wave sleep [3]. Instead, the risk is strictly additive, following a severe linear dose-response curve where cognitive errors escalate dramatically as independent behavioral stressors aggregate (Figure 2d).In a biological system like ourselves, this architecture is likely bidirectional and recursive-distress fragments sleep, which subsequently saps the motivation for physical activity, trapping the individual in a feedback loop that heightens cognitive failure rates. Consequently, optimizing daily cognitive function requires moving beyond single-variable interventions. Because these behavioral burdens accumulate linearly rather than interactively, clinical and lifestyle therapies must comprehensively target this triad, prioritizing sleep preservation and distress reduction. Future longitudinal investigations utilizing objective actigraphy and high-density EEG will be critical to fully untangle these recursive dynamics and map the long-term stability of these neural representations.

## Declarations

### Ethical approval

This study was conducted in accordance with the ethical principles of the Declaration of Helsinki. Informed digital consent was obtained from all participants prior to data collection.

### Author contributions

AS: study conception, data collection, data analysis, interpretation, drafting, and revision of the manuscript.

### Data availability

Data supporting this study are available from the corresponding author upon request.

